# mSOD1-NSG mice: a new in vivo model to test human T cells in ALS

**DOI:** 10.1101/2021.09.27.461982

**Authors:** David J Graber, Marie-Louise Sentman, W. James Cook, Charles L Sentman

## Abstract

Amyotrophic lateral sclerosis (ALS) is a progressive neurodegenerative disease with unclear etiology and few treatment options. Engineering of human cells for therapy has great potential to provide a means to create long-lived therapeutics that can respond to local signals within tissues. One challenge for development of human cell therapeutics is to have disease models that do not reject transplanted human cells. G93A mutant superoxide dismutase-1 (mSOD1) transgenic mouse model is a robust disease model for ALS, with development of local inflammation, activation of microglia and astrocytes, motor neuron death, and development of paralysis. To create a mouse model for ALS that permits the transplantation of human T cells without immune-mediate rejection, we bred the G93A transgene onto the NOD-SCID-IL-2Rγ-deficient (NSG) mouse model to create mSOD1-NSG mice. We report that mSOD1-NSG mice develop a progressive ALS-like disease with microgliosis, astrogliosis, and paralysis development with an average onset at 11 weeks and end-stage at 14 weeks. Transplanted human T regulatory cells survive in these mice for at least 60 days. This mSOD1-NSG mouse model for ALS can be used to test human T cell-based therapeutics, and it may be helpful to test any human cell-based therapy for ALS.

**Significance Statement:** G93A SOD1 transgenic NSG mice develop a progressive ALS-like disease and are an in vivo model for testing human T cell-based therapies for ALS.

## Introduction

Amyotrophic lateral sclerosis (ALS) is a progressive neurodegenerative disease with unclear etiology and there are few treatment options that only have minimal effect. The chronic degeneration of motor neurons leads to an average survival of less than five years after diagnosis (1). There are approximately 6,000 new cases of ALS in the United States each year. About 10% of ALS cases are familial cases, and 90% are spontaneous (2). A number of genes have been linked to ALS, and one of the most studied is superoxide dismutase-1 (SOD1). Mutations in SOD1 have been shown to be linked to familial cases of ALS and to occur in spontaneous ALS cases as well (2–4). Several of these mutations have been used to create mutated SOD1 transgenic mice, rats, and pigs that recapitulate many of the features of human ALS (5–7). A G93A mutation in SOD1 is associated with human ALS and transgenic mice that express this mutated human protein develop a progressive motor neuron disease, inflammation in the spinal cord, and these mice progressively develop limb paralysis (8–10). Depending on the genetic background on which the mutant SOD1 is expressed, the progression of the disease can be slower or faster, with B6/SJL mixed strain have an average lifespan of 129 days, while a B6 background strain lives around 144-161 days, a SJL strain lives around 119 days, and the 129 strain lives around 125 days (10, 11). Other SOD1 mutations such as G85R, G37R, and D90A also result in motor neuron degeneration and paralysis in transgenic mice, but the disease proceeds at a slower rate and/or later onset with these SOD1 mutations.

There are no therapeutics available that can stop the progression of ALS and few that can slow the rate of disease. One challenge for drugs, biologics, and other therapies is to access the sites of disease within the brain and spinal cord due to the blood-brain barrier. There are several ways to get therapies passed the blood-brain barrier through drug design, by bypassing the barrier through direct CNS injection, or using cell-based therapies for drug delivery. Cells of the immune system readily access the CNS as a part of their physiological trafficking. Although once thought to be immune privileged, data have shown the presence of T cells, B cells, macrophages and other immune cells accessing the CNS under physiological and pathological situations (12, 13). The presence of inflammation within the CNS increases different types of immune cells that can help to heal or exacerbate the disease process. ALS is associated with an increase in inflammatory cells and cytokines within the spinal cord in human patients and in mutant SOD1 transgenic mouse models (14–16).

It has been shown that as ALS progresses there are changes in the peripheral immune system. T regulatory cells (Tregs) are a subset of immune T cells that limits inflammation, autoimmunity and helps to maintain homeostasis. Tregs can be isolated and expanded in vitro, genetically engineered with defined specificities, and they are being explored as adoptive cell therapy for autoimmune diseases and transplantation (17–19). As ALS disease progresses, the number of Tregs decreases and a low number of Tregs is associated with worse outcomes in patients (20, 21). As an approach to reduce inflammation within the spinal cord, it is possible to isolate, expand and infuse Tregs into ALS patients. Data from a small phase I clinical trial by Stanley Appel and his team at Baylor showed the feasibility of this approach and demonstrated initial safety (22). Other cell-based therapy approaches involve injected human mesenchymal stem cells or iPSC-derived cells directly into the spinal cord (2, 23, 24). One challenge with studying these approaches and their further development is that immune intact animal models can be associated with rejection of human cells. Depending on the cell dose and location, the rejection of xenogeneic tissue occurs at varying rates. In this study, we developed an immune deficient mouse model with the G93A human SOD1 transgene that allows persistence of transplanted human T lymphocytes. The mSOD1-NSG mice develop a rapid and robust ALS-like disease and allows the survival of human Tregs in vivo for many weeks.

## Material and methods

### Animals

NOD.Cg-Prkdc^scid^ IL2rγ^tm1Wjl^ (NSG) mice were purchased from the Dartmouth Mouse Modeling Shared Resource (Lebanon, NH). SOD1^G93A^ B6 mice (stock # 004435), SOD1^G93A^ B6SJL mice (stock # 002726), and Rag2-deficient B6 mice (stock # 008449) were purchased from the Jackson Laboratory (Bar Harbor, ME). Mouse experiments and procedures were ethically conducted under the approval of Dartmouth College’s Institution Animal Care and Use Committee. Mice were bred and colonies maintained at Dartmouth’s Center for Comparative Medicine and Research facility.

Male SOD1^G93A^ B6 mice were bred with female NSG mice to develop SOD1^G93A^ NSG mice, which we refer to as mSOD1-NSG mice (Figure 1). Tail tissue samples were collected after weaning and used to determine human SOD1 genotype and transgene copy number according to the Jackson Laboratory (Bar Harbor, ME) recommended primers and procedures, and controls included were DNA from an initial SOD1^G93A^ B6 mouse (positive) and NSG mice (negative) in each assay. A value of positive control (C_t_^hSOD1^ - C_t_^ApoB^) – sample (Ct^hSOD1^ - Ct^ApoB^) of greater than 0.7 was considered a low copy number. To determine which mice in the backcrossed generations were deficient in lymphocytes, 0.2 mL of blood samples were collected and analyzed by flow cytometry for absence of CD3+CD4+ cells prior to breeding age. All male SOD1 transgenic mice were typed in BC1, BC2, and BC3 generations to select mice for breeding to NSG female mice. mSOD1-NSG mice were monitored for weight and assessed for limb paralysis three-times-per-week starting at 7 weeks of age. Peak weight was determined using a moving average of weight over a seven-day span. Mice that lost greater than 15% of peak weight were considered at disease endpoint and sacrificed.

**Figure 1.**
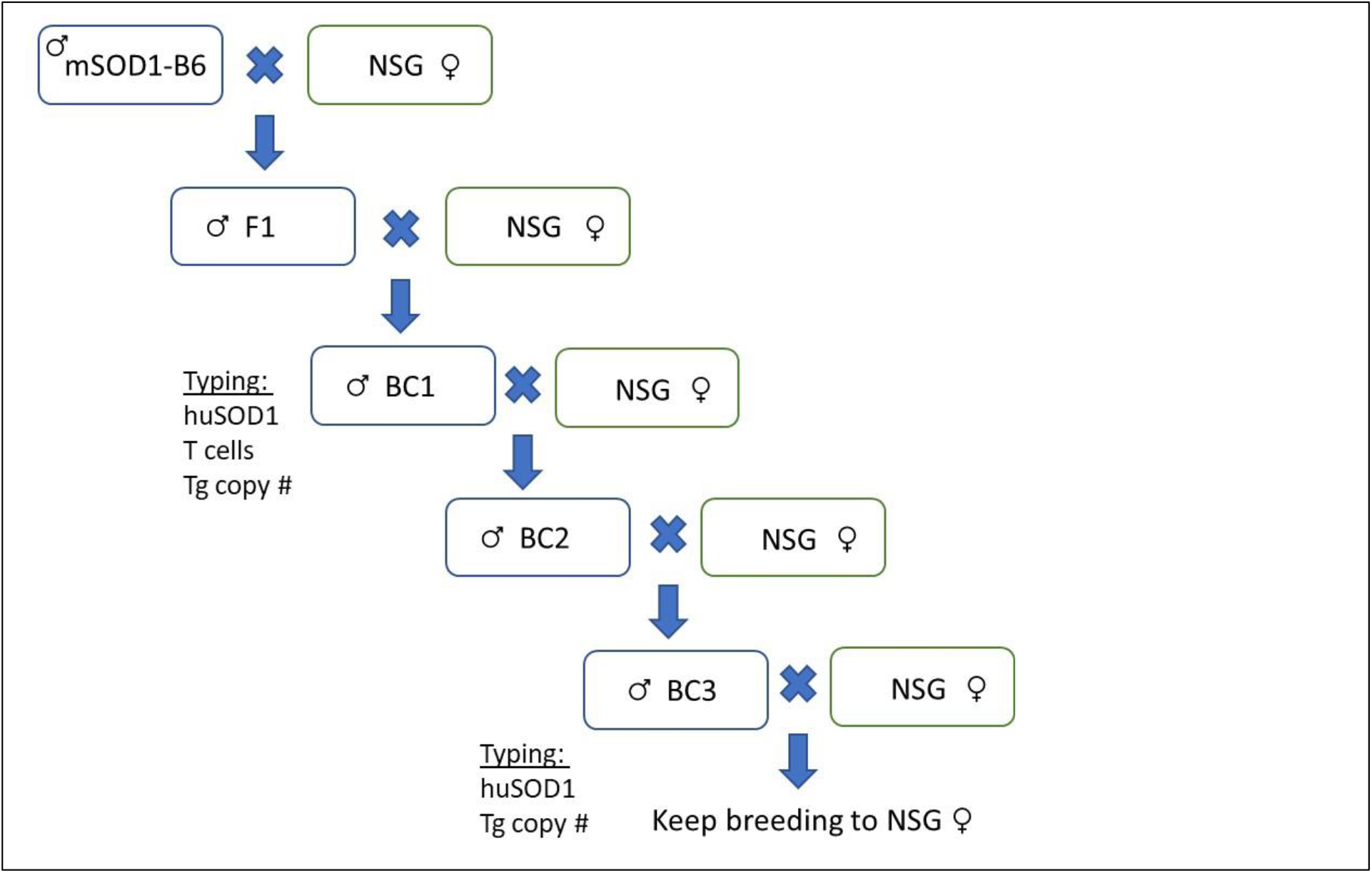
Breeding scheme to create mSOD1-NSG mice. G93A mSOD1-B6 mice were bred to NSG mice, and offspring backcrossed to NSG mice. BC1 male mice were typed for huSOD1 transgene, white coat color, lack of blood lymphocytes, and transgene copy #. All male offspring used for breeding from BC2 and BC3 generations were confirmed for SOD1 transgene and T lymphocyte deficiency. These mice are maintained as G93A SOD1-transgene heterozygous mice by breeding with NSG female mice. Offspring are typed for the SOD1 transgene and copy #.

mSOD1-NSG and two additional mouse models were used to evaluate survival and persistence of human T cells in vivo: (1) Rag2-deficient B6 mice and (2) SOD1^G93A^ B6SJL mice. The SOD1^G93A^ B6SJL mice (only white coat fur mice were used) received6 Gy whole body sublethal irradiation from a cesium-137 irradiator (J.L. Shepherd & Associates, San Fernando, CA) one day prior to human cell injection. Human Tregs were injected i.v. in the tail vein with mice in a restrainer chamber. When noted, human IL-2 was injected at 40,000 U/mouse i.p. on days 2, 4, and 6 after injection of cells. Monitoring of luciferase-expressing human cells in vivo was performed by injecting mice i.p. with 3.75 mg of luciferin (GoldBio) diluted in PBS. Mice were anesthetized with isofluorane and imaged six-to-eight minutes later using an IVIS system (Xenogen). Rag2-deficient B6 mice were shaved prior to first imaging. Images were analyzed using Living Image software (PerkinElmer) with the same exposure time and photon scale.

### Treg isolation and expansion

Human Tregs were enriched or purified from PBMCs collected from anonymous donors immediately following plasmapheresis (Blood Donor Program, Dartmouth-Hitchcock Hospital, Lebanon, NH). CD4+ cells were isolated by negative-selection using MOJOSORT™ Human CD4 T Cell Isolation Kit (Biolegend) and a EASYSEP™ Magnet (StemCell) according to manufacturer’s instructions. Then, CD25hi cells were enriched from these CD4+-isolated cells by positive-selection using anti-human CD25 MicroBeads II (Miltenyi) and MS Columns with MiniMACS™ Separator magnet (Miltenyi). The enriched CD4+CD25hi cells were cultured in 24-well non-tissue culture plates at 1 x 10^6^ cells/mL in X-Vivo-15 (Lonza) supplemented with 10% heat-inactivated human AB serum (Sigma). Cell incubation condition was humidified 37°% CO_2_. Cells were stimulated with 25 μL/1 x 10^6^ cells of ImmunoCult™ Human CD3/CD28 T Cell Activator (StemCell) on days 0 and 9 in culture. Culture media was supplemented with 500 U/mL human IL-2 (Tecin from Roche, kindly provided by the NIH) starting on culture day 2. Cultured cells were transferred to 25cm^2^ tissue culture-treated flasks on day 5. Cells were transduced over two days with a retroviral construct containing a PpyRE9 firefly luciferase fused to eGFP genes on day 10 and 11 in culture. To transduce Tregs, 24-well non-tissue culture plate wells were pre-coated with RetroNectin® (Takara Bio USA, Inc.) according to manufacturer’s instructions and then day 10 cultured Tregs were added at 0.3 x 10^6^ cells/well in 0.3mL of cell culture media. Retroviral supernatant was added at 0.7mL/well and plates were centrifuged at 1500 rcf at 30°C for 1h, and then incubated overnight. On the next day, 0.5mL culture supernatant was replaced with 0.5mL retroviral supernatant with 500 U/mL IL-2 and cells were re-centrifuged and then incubated overnight. The next day cells were transferred to 25cm^2^ tissue culture-treated flasks at 1 x 10^6^ cells/mL. On day 13 of culture, a sample of cells was evaluated for luciferase transduction efficiency by measuring the percentage of the co-transgene eGFP on cells by flow cytometry. Cells were cultured with fresh cell culture media added every two days until day 17. Cells were evaluated by flow cytometry prior to injection into mice and were 92.6% ± 2.2% CD4-positve with increased staining for FoxP3.

When noted, human Tregs were further purified following the positive-selection step using anti-human CD25 MicroBeads II and MS Columns with MiniMACS™ Separator magnet. The CD4+CD25hi cells were then additionally labeled with CD4-FITC (Biolegend) and CD127-APC (Biolegend) and sorted for CD4+CD25hiCD127lo using FACSAria III Cell Sorter carried out in DartLab, the Immune Monitoring and Flow Cytometry Shared Resource at the Norris Cotton Cancer Center at Dartmouth. CD25hiCD127lo purified Tregs were cultured and transduced as described above except 500 U/mL IL-2 was added to the culture medium starting on day 0. On day 17 of culture cells were evaluated by flow cytometry prior to injection into mice and were 99.4% ± 0.1% CD4-positve with increased staining for FoxP3. These cells are referred to as FACS purified Tregs or FP Tregs.

### RNA extraction, cDNA preparation, real time qPCR

Whole spinal cords were collected from mSOD1-NSG at indicated stages of disease and stored in RNA Protect (Qiagen). Spinal cords were removed from RNA Protect solution and RNA was extracted using TRIzol Reagent (Life Technologies, Carlsbad, CA, USA). The eluted RNA was quantified by NanoSpec spectrophotometry, and 2 μg was reverse transcribed using qScript cDNA SuperMix (Quanta Biosciences, Gaithersburg, MD, USA). Mouse oligonucleotide primer sets for β-Actin, CCL2, CCL4, Nox2, and IL-1β were described previously (25). Mouse TNF-α primers were Fwd-ACCACGCTCTTCTGTCTA and Rev-GAAGATGATCTGAGTGTGAGG. Quantitative real-time PCR was performed using PerfeCTa SYBR Green FastMix without ROX (Quanta Biosciences), 10 ng of cDNA, and 300 nM of a RT-PCR primer sets (IDT, San Jose, CA, USA). Settings for analysis using a C-1000 CFX96 machine (Bio-Rad) were as follows: initial denaturation (95°C/2 min) was followed by 45 cycles of denaturation (95°C/10 s) and primer annealing (60°C/30 s). A melt curve was performed on all samples for quality control. Data was quantified by the 2(-ΔΔCt) method using β-actin as an internal control reference mRNA.

### Flow Cytometry

Flow cytometry antibodies: mouse CD3-APC (Biolegend), mouse CD4-PE (Biolegend), human CD4-APC (Biolegend), human FoxP3-APC (eBioscience). For FoxP3 staining cells were permeabilized using Foxp3 / Transcription Factor Staining Buffer Set (eBioscience). Cells were analyzed for eGFP transduction or staining antibodies with a C6 Accuri flow cytometer (BD Biosciences).

### Immunohistochemistry

Spinal cord sections were cut into 3-4 mm long segments, fixed in formalin overnight, and paraffin-embedded by Pathology Shared Resource (Dartmouth-Hitchcock hospital). Four-micron thick transverse-sections on microscope slides were deparaffinized and antigen retrieval was performed in 10 mM citric acid (pH 6.0) + 0.5% Tween 20 for 3 min in an Insignia pressure cooker. IHC was performed in 0.1 M PBS with 5% goat serum. Sections were treated overnight at 4°C with primary antibodies: rabbit anti-Iba1 (1/1200, Wako), chicken anti-GFAP (1/1500, Biolegend), mouse anti-NeuN (1/100, EMD Millipore). Sections were treated for 2 h at RT with secondary goat antibodies (1/400, Invitrogen): anti-rabbit-Al488, anti-chicken-Al649, and anti-mouse Al568. Tissue sections without primary antibody were used to control for background staining. Sections were cover slipped using SlowFade Antifade Reagent with DAPI (Thermo Fisher Sci). Spinal cord sections were imaged using Olympus IX-73 Inverted Fluorescence Microscope with a 20X objective and an Olympus DP73 CCD camera using consistent exposure times. Images were merged using Image J 1.53E software (NIH).

### Statistics

Statistical analysis was conducted using GraphPad Prism Software (La Jolla, CA, USA). Significant differences were defined as *P < 0.05,** P < 0.01, and *** P < 0.001. Changes in age of peak weight, paralysis onset, and disease endpoint were compared among male and female mSOD1-NSG mice using Kaplan-Meier survival statistics (log rank sum test) and Cox’s F test comparison. One-way ANOVA followed by Dunnett’s test was used to compare mRNA in mSOD1-NSG mice at different disease stages relative to non-transgenic littermate controls. One-tailed student t-Test was used to compare in vivo bioluminescence in Treg-injected mice versus background signal in vehicle-injected mice. One-way ANOVA followed by Tukey test was used to compare in vivo bioluminescence in PBS and IL-2 injected mice.

## Results and Discussion

### Generation of mSOD1-NSG mice

In order to create an in vivo model for ALS that would allow the incorporation of human therapeutic cells, we chose to breed G93A SOD1 transgenic C57BL/6 mice to NOD-SCID-IL2Rγ-deficient (NSG) mice. NSG mice not only lack T, B, and NK cells, they also have a mutation in SIRP1-alpha so that they do not recognize CD47 on human cells (26, 27). This protects human cells from macrophage-mediated cell killing in NSG mice. Male G93A SOD1 transgenic mice were bred to female NSG mice, and the resulting male F1 progeny were backcrossed to NSG mice. These first-generation backcrossed offspring (BC1) were expected to be 75% NSG background and 50% having the G93A transgene and 50% having the SCID mutation. The IL-2Rγ chain is found on the x chromosome, so by choosing male offspring from female NSG mice, all of the male mice will have the defective IL-2Rγ chain. The BC1 mice were typed for the human SOD1 transgene by PCR, then blood was taken to type for the absence of lymphocytes. Mice with the SCID mutation will have very few T or B cells, while the I L-2Rγ-deficiency will result in the absence of NK cells too. About 25% of the BC1 mice were both G93A SOD1 transgenic and had no T cells in the blood. We chose the male mice to bred to NSG mice to create the BC2 progeny. The male mice were typed for the G93A SOD1 transgene and for a lack of blood T cells to confirm their phenotype. Mice were also selected for the white coat color found in the NSG strain because this trait makes in vivo imaging by luminescence about five times more sensitive compared to mice with a dark coat color. Male mice were selected and bred to NSG female mice to create the BC3 progeny. All SOD1 transgenic BC3 male mice had a white coat color and lacked blood T cells. The BC3 mice were expected to be ~94% of the NSG background, and 50% had the SOD1 transgene. We took BC3 male mice and bred them to NSG females to create an expanded cohort of BC4 mice for experimental use and analysis of the ALS phenotype. An overview of the breeding scheme is shown in Figure 1. From this point forward, the mSOD1-NSG transgenic mice were also typed for transgenic copy number as well. It is known that a loss of copy number results in a delay in the disease onset (28). We have found that less than 1 in 30 progeny have a reduced copy number from the initial founder SOD1^G93A^ B6 mouse, and these mice were removed from experiments. All breeding SOD1 transgenic mice have retained a high copy number, which is tested prior to breeding. We have continued to take male offspring typed for the SOD1 transgene and high transgene copy number for breeding of more than 8 generations. Note that the onset, paralysis, and survival curves have not changed since we started studying these mice in BC4 through to the BC8 generation. We refer to this mouse strain as mSOD1-NSG mice.

### Rapid ALS-like disease development in mSOD1-NSG mice

G93A transgenic mice have a well-documented disease course, although the timing of onset, paralysis, and endpoint varies for different background strains (11). Although inflammation is a key aspect of the disease process, T cell-deficient G93A SOD1 transgenic B6 mice, that lack functional T cells, have an accelerated disease course compared to immune intact G93A SOD1 transgenic B6 mice (14, 29). Body weight is a non-subjective measure of overall animal health and SOD1 transgenic mice reach a peak weight and then lose weight until the endpoint is reached, whereas non-transgenic mice continue to gain weight (11). mSOD1-NSG mice were weighed three times-per-week and average weight was calculated over 7 days, and this average weight was used to calculate the maximum (peak weight) for each mouse. For ethical considerations, we chose a decrease of > 15% of peak weight as our endpoint. We have also used a loss of > 20% body weight in some experiments, and the results were similar with a longer survival by about 2 days. The onset of paralysis was determined by testing grip on a wire cage after each weight measurement and a lack of grip on any paw was noted (usually a hind paw) as the start of paralysis. Almost all mice developed complete paralysis in at least one limb before reaching the 15% weight loss endpoint.

The data in figure 2 show onset (peak weight), limb paralysis development, and overall survival (age at endpoint). Onset of weight loss occurred at an average age of 11.1 ±1.2 (±SD) weeks. Limb paralysis developed at 13.9 ±1.2 weeks on average. Overall survival was averaged 14.1±1.2 weeks, although a few mice reached endpoint at 12 weeks and others survived more than 16 weeks. There was no significant difference between male and female mice in these readouts, although the overall survival and limb paralysis showed greater variance with female mice. Thus, a greater number of animals are required to show statistical differences when female mice are included in studies, whereas the male mice will have less variance within groups. These data show that the mSOD1-NSG strain has a similar disease course as other G93A SOD1 transgenic mice, but that the disease course is faster than G93A transgenic SOD1 mice on the B6 background, which have an average onset around 12.5 weeks and reach end stage around 20 weeks. This suggests that treatment approaches in mSOD1-NSG mice need to be started earlier than in immune intact mSOD1-B6 mice and it also supports the idea that an intact immune response helps to delay the disease process.

**Figure 2.**
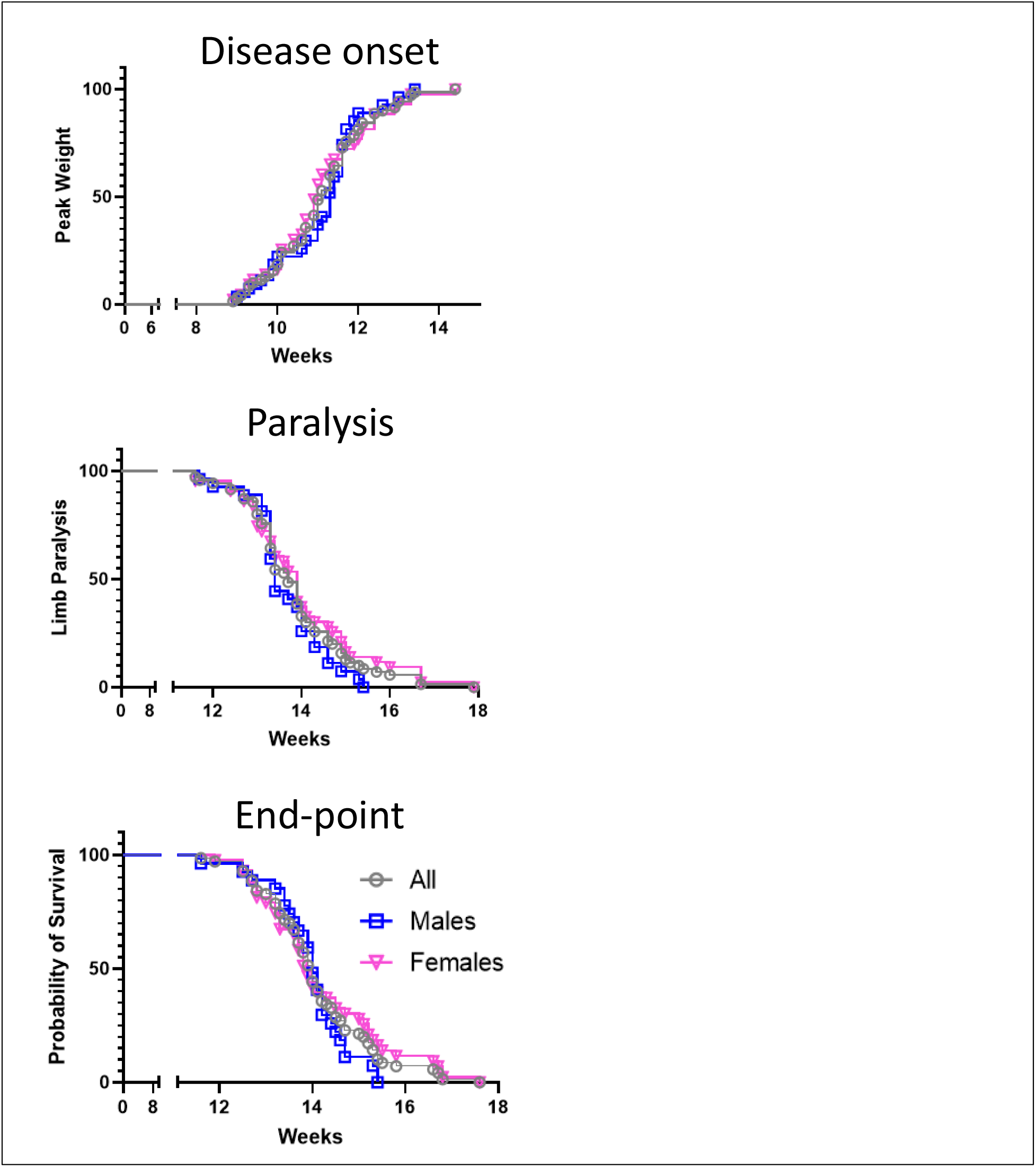
mSOD1-NSG onset, paralysis, and end-point. The phenotype of mSOD1-NSG mice as a function of age (in weeks). The data are shown for all mice combined (grey circles, n = 70), males (blue squares, n = 27), and female (pink triangles, n = 43). There were no statistical differences between the sexes on these readouts in the mSOD1-NSG mice by Log-rank test.

### An increase in microglia and astrocyte markers in the spinal cord at end-stage

A hallmark of ALS is the activation and increase in microglia and astrocytes within the spinal cord (6, 9, 30, 31). Of note, due to the NOD genetic background, macrophages and microglia in NSG mice have a mutation in SIRP-1α and a defective IL-2Rγ protein, which affects type 1 IL-4 receptors and IL-15 receptors. We examined microglia, astrocytes and neurons in the spinal cord at end stage. Figure 3 shows typical staining for Iba1 (microglia), GFAP (astrocytes), NeuN (neurons), and DAPI (cell nuclei) in spinal cord sections from end stage disease and a non-transgenic littermate control. There is an increase in the intensity of Iba1 (microglia) and GFAP (astrocytes) expression and noticeable decrease in large soma NeuN staining (neurons) in the mSOD1-NSG transgenic spinal cord samples.

**Figure 3.**
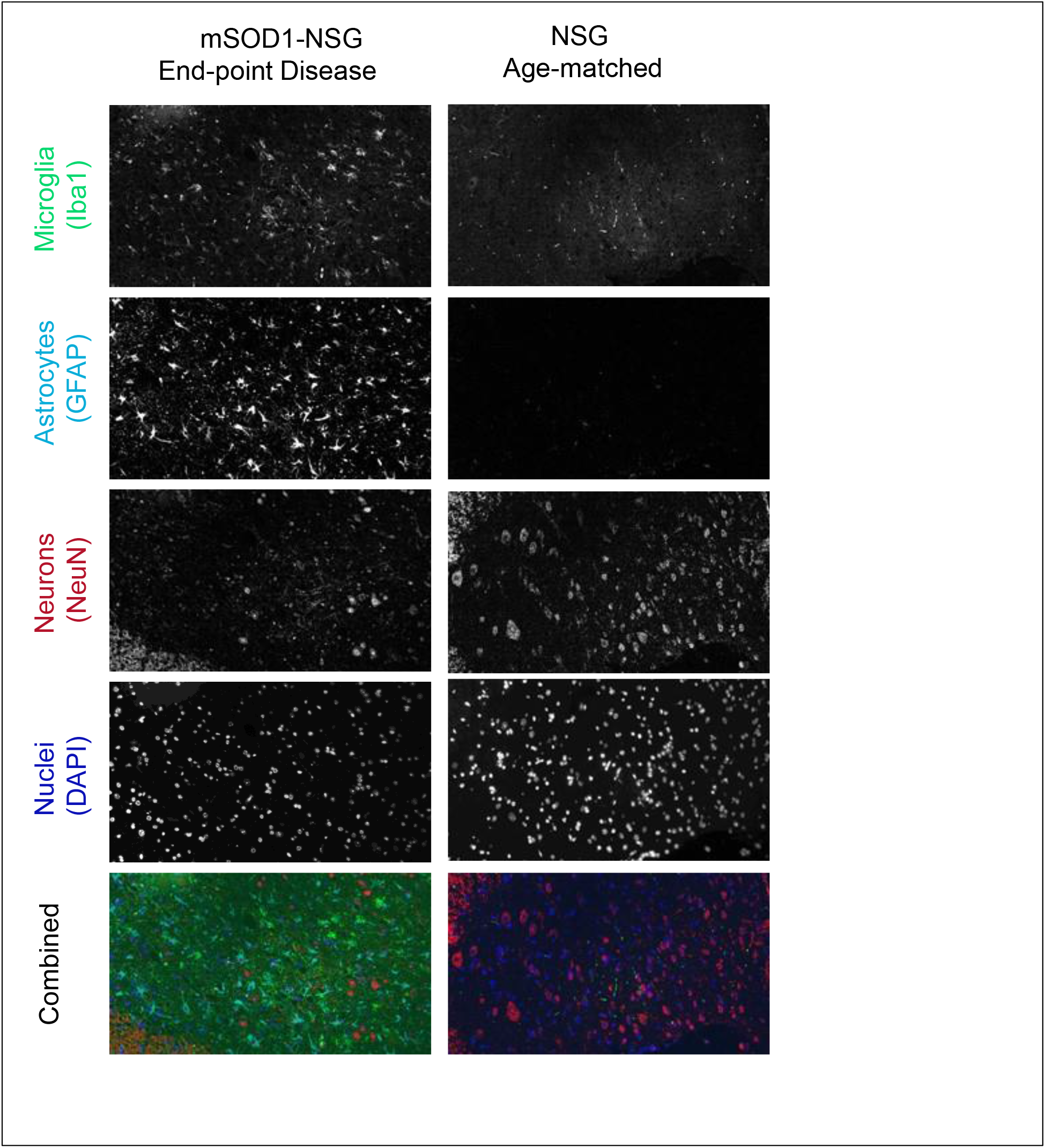
Microglia, astrocytes, and neurons in spinal cord of mSOD1-mice at end-point. Spinal cords from mSOD1-NSG mice at disease endpoint and littermate NSG controls were taken, fixed, cut, and sections stained for Iba1 (green), GFAP (cyan), NeuN (red), and DAPI (blue). Representative data are shown.

### Pro-inflammatory cytokines are found in the spinal cords of mSOD1-NSG mice and increase as disease progresses

The role of inflammation in neurodegeneration is controversial, but as disease progresses, local neuroinflammation is increased(30, 32–34). To examine inflammatory mediator expression within the spinal cord of mSOD1-NSG mice at early and late stages of disease, we removed spinal cord, isolated RNA, made cDNA and determined the expression of specific cytokines, chemokines, and other mediators by qPCR (Figure 4). Data are shown as fold change compared to non-transgenic littermates. Each symbol represents a single individual. We analyzed three different stages of the disease – preclinical (8.1 to 9.6 weeks), onset (10.1 weeks to 12.0 weeks), and endpoint (> 15% weight loss). At an age close to typical peak weight, there is an increase in CCL2, CCL4, IL-1beta, and TNF-alpha, with greater amounts at end stage disease. Many of these cytokines are increased 5-fold compared to the littermate controls by end stage. NOX-2, an enzyme involved in production of oxygen free radicals, is elevated 4.5-fold at the clinical stage of disease. It is important to note that the lack of shared cytokine receptor signaling via IL-2Rγ did not prevent the development of spinal neuroinflammation in these mice.

**Figure 4.**
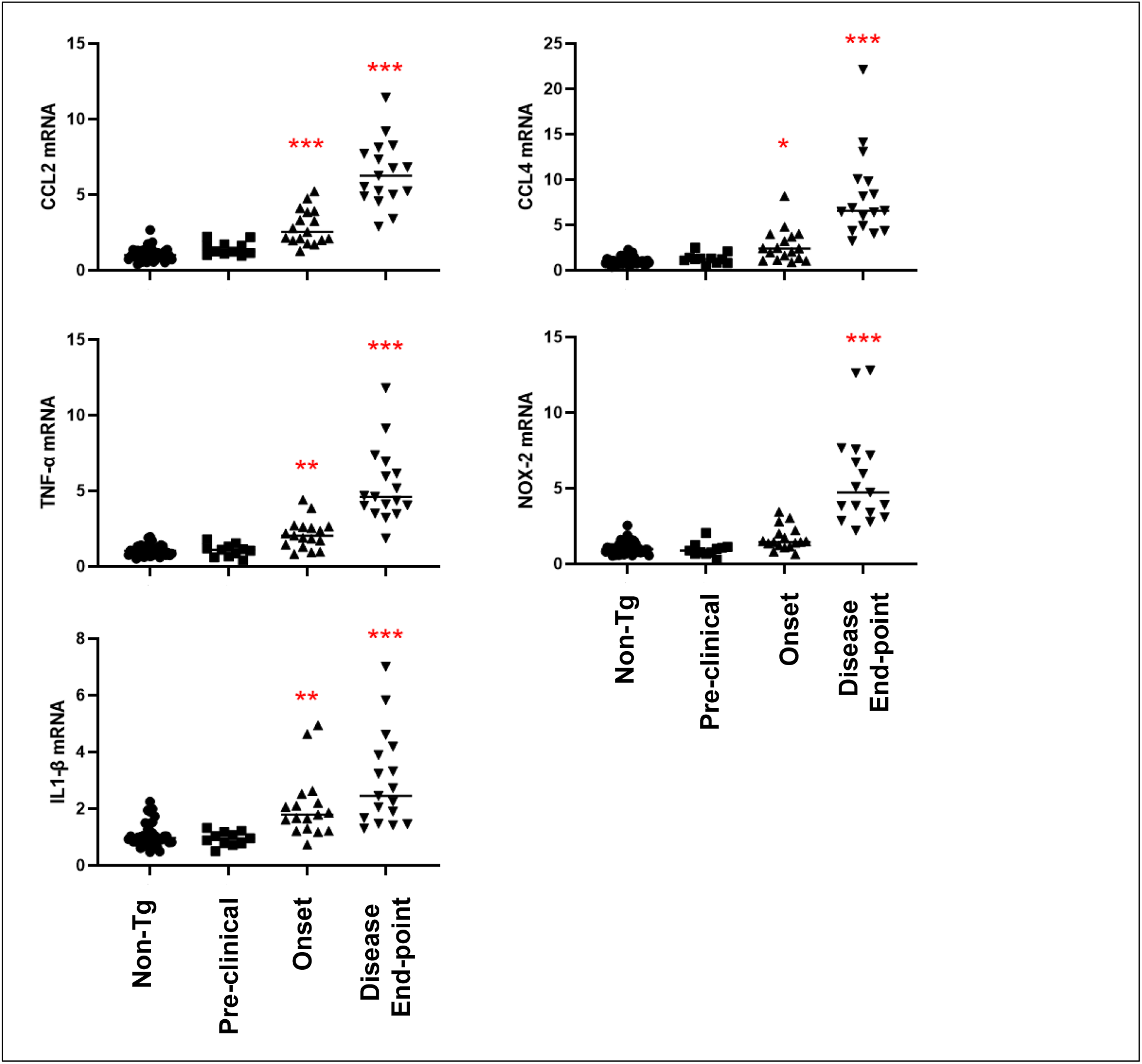
Inflammation increases as disease progresses in mSOD1-NSG mice. Spinal cords from mSOD1-NSG mice or littermate NSG controls (n = 40) were taken at preclinical (8.1 to 9.6 weeks, n = 11), near disease onset (10.1 to 12.0 weeks, n = 17) and at endpoint (>15% weight loss, n = 17). Tissue was dissected and processed for RNA, then cDNA. Real-time qPCR was used to determine the expression of CCL2, CCL4, TNF-α, NOX-2 and IL-1β compared to β-actin. Relative expression compared to the average of the NSG littermate controls was determined. Each point is an individual sample and the group average is indicated by a line. * p<0.05, ** p<0.01, *** p<0.001 by a Dunnett’s test.

### Human Tregs are detected in mSOD1-NSG mice 60 days post IV transfer

One of the goals of this study was to create a mouse model for ALS where human cells could survive in vivo without immune-mediated rejection. To study the tracking of injected human Tregs, we transduced sorted, expanded human Tregs with luciferase to allow in vivo tracking using an IVIS system. To allow for longer term assessment of cell survival in vivo in the absence of disease development, we used the non-transgenic littermates (no G93A mSOD1) to study persistence of human Tregs. As shown in Figure 5, the human Tregs that were enriched, expanded in vitro, and injected IV were able to be detected within the mice for up to 9 weeks following transfer. The numbers of cells decrease over the first 30 days, then appear to stabilize in the mice. The injected Tregs were initially spread throughout the body, as expected after IV injection, but at later time points, Tregs were found in primarily in the upper body and head region. However, human Tregs injected into B6-Rag2 deficient mice were rapidly lost. Similarly, low dose ionizing irradiation can be used to decrease WBCs and create a niche for transplanted lymphocytes to expand. However, human Tregs were completely gone within 2 days after injection into mSOD1 B6/SJL mice pretreated with sublethal whole body irradiation.

**Figure 5.**
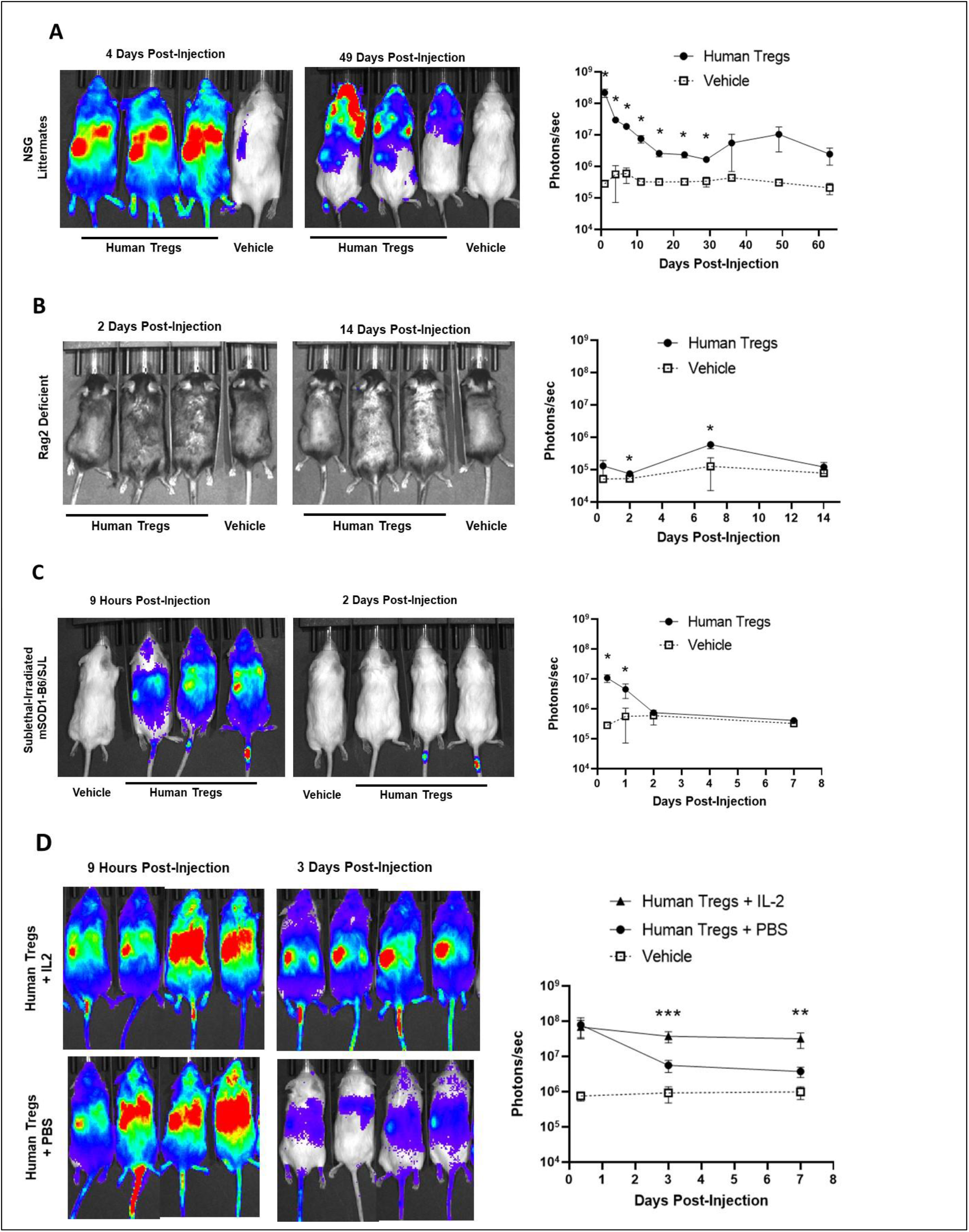
Persistence of human cells in vivo. Mice were injected IV with luciferase expressing human Tregs. Cells were tracked using Xenogen IVIS at various times after injection, as indicated. Representative images are shown on the left, and average total signal detected for each group (photons/sec) are graphed on the right. Expanded human Tregs were injected into (A) non-transgenic littermates of mSOD1-NSG mice (n = 4), (B) Rag1-deficient B6 mice (n = 3), (C) sub-lethally irradiated mSOD1 B6/SJL mice (n = 5), or (D) non-transgenic NSG littermates with (n = 4) or without (n = 4) IL-2 IP on days 2, 4 and 6. Error bars show StDev. * p<0.05, ** p<0.01, *** p<0.001 by one-tailed student t-test (panels A-C) or ANOVA followed by Tukey test (panel D).

Human Tregs are cultured in high amounts of IL-2 and NSG mice do not have an endogenous source of IL-2. This dependence on IL-2 may result in a loss of human Tregs in vivo, and IL-2 has been given to clinical patients injected with in vitro expanded Tregs (18, 22, 35). To test whether additional IL2 treatment would promote the persistence of Tregs in these mSOD1-NSG mice, mice were given injections of IL-2 i.p. on days 2, 4 and 6 after Treg injection. For these experiments, we used FACS purified (FP) Tregs, which were further purified by cell sorting before in vitro expansion. These FP Tregs were 99.4% CD4+ with high FoxP3 expression after 17 days in culture. Mice that were injected with human Tregs in combination with IL-2 treatment maintained Tregs well over the first week, whereas mice given Tregs without IL-2 had much lower signal during the first week after injection suggesting that fewer cells survived or proliferated. We have found that the human Tregs can be found several weeks after treatment in the spleen and in spinal cord of mSOD1-NSG mice even though no additional IL-2 is given after day 6 (data not shown). Of note, conventional effector human CD4 and CD8 cells can cause xenogeneic GvHD in NSG mice when given at large cell doses. However, no signs of GvHD (hair loss) were observed in our studies using human Tregs. Overall, the data show that human lymphocytes survive in mSOD1-NSG mice for many weeks and initial IL-2 support limits the initial loss of cells.

## Acknowledgements

The authors wish to thank the staff of the Center for Comparative Medicine Research for support for the animal work. The staff of the Department of Pathology at Dartmouth Hitchcock Medical Center for tissue preparation. The authors wish to thank Gary Ward for cell sorting. Cell sorting was carried out in DartLab, the Immune Monitoring and Flow Cytometry Shared Resource at the Norris Cotton Cancer Center at Dartmouth, with NCI Cancer Center Support Grant 5P30 CA023108-41. DNA sequencing was carried out at the Geisel School of Medicine at Dartmouth in the Genomics Shared Resource, which was established by equipment grants from the NIH and NSF and is supported in part by a Cancer Center Core Grant (P30 CA023108) from the National Cancer Institute. The authors also thank the National Cancer Institute Biological Resource Branch for recombinant human IL-2.

## Author contributions

DJG and CLS designed the project; DJG, MLS, WJC acquired the data; DJG, MLS, WJC, CLS analyzed and interpreted the data; and DJG and CLS wrote the paper.

## Notes

**Competing Interest Statement:** The authors (DJG, WJC, and CLS) have a pending patent application for chimeric antigen receptors (CARs) and the use of CAR T-regulatory cells as therapy for neurodegenerative diseases. These interests are managed through the policies of Dartmouth College.

**Funding:** This work was supported in part by grants from the National Institutes of Health (NS102556 and NS117895) and funds from the Center for Synthetic Immunity, the Geisel School of Medicine at Dartmouth.

### Competing Interest Statement

The authors (DJG, WJC, and CLS) have a pending patent application for chimeric antigen receptors (CARs) and the use of CAR T-regulatory cells as therapy for neurodegenerative diseases. These interests are managed through the policies of Dartmouth College.

